# A Scalable and Robust Workflow for Cost-Effective Post-Translational Modifications Profiling by Chemical Proteomics

**DOI:** 10.64898/2026.08.17.745240

**Authors:** Liu Zang, Jonas Grandke, Jana Richter, Pavel Kielkowski

## Abstract

Mass spectrometry-based chemical proteomics is a powerful method to analyze proteins labelled by small molecules to identify protein targets of active compounds and to profile protein post-translational modifications. The throughput and high protein input for chemical proteomics workflows has been often a limiting factor for application of the technology for specialized and difficult to culture cell lines. The high protein input was necessary to gain significant difference of noise to signal ratio in proteomics readout. Here, we describe a general chemical proteomics workflow, which is performed in 96-well plate and necessitate only 25 μg of protein input to profile post-translationally modified proteins including abundant *O*-GlcNAcylated proteins as well as low abundant AMPylated proteins. The workflow integrates advances in Cu(I)-catalyzed azide-alkyne cycloaddition to minimize chemical side-reactivity of the ‘click reaction’ and data-independent acquisition mode during LC-MS/MS measurement. An iterative optimization of protein clean-up on carboxylate-coated paramagnetic beads led to significant saving of the beads usage and lowers the unspecific protein background that resulted in sensitivity gain.

## INTRODUCTION

Chemical proteomics is an indispensable tool for identification of protein targets of active small molecules including pharmaceutically active compounds and protein post-translational modification (PTM) mimics.^1–3^ Although the mass spectrometry-based proteomics has undergone rapid technological development that has significantly improved sensitivity and depth of the analyzable proteome, it is not yet possible to detect and quantify all low abundant protein modifications collectively known as proteoforms; regardless of their origin that might arise from a chemical reactivity or metabolic activity.^4–6^ Hence, the chemical proteomics takes advantage of small molecule probes or reporters, which can facilitate the enrichment of the proteoforms and so their detection by mass spectrometry-based proteomics. In the chemical proteomics workflow, a small compound probe containing a chemical handle, typically a terminal alkyne or azide, is added to a cell culture media to allow its interaction with proteins and after the cell lysis, the probe labelled proteins can be reacted by ‘click chemistry’ with a complementary affinity tag to enable their enrichment.^7,8^ The enriched proteins are subsequently cleaved into peptides by trypsin, which are resolved by LC-MS/MS. The acquired mass spectra are then searched against the protein database to identify the enriched proteins. In order to distinguish the probe-labelled proteins from the background composed of proteins nonspecifically interacting with the solid support used for the enrichment, it is necessary to compare a probe treated proteins with a control, which does not contain the probe. In recent years, several chemical proteomic workflows have been developed to increase the throughput of the method including FAIMS-SP3^9^ and approaches combined with a chemical labelling of proteins for their quantification such as tandem mass tags (TMT).^10^ The main difference to a traditional approaches is the streamlined protein clean-up after the ‘click reaction’ previously using the protein precipitation which is usually the most time and labor intensive step.^11^ Indeed, overall reduction of the hands-on time and number of protein transfer steps between tubes is highly desired. Furthermore, to enable the practical application of chemical proteomics for studies of specialized and difficult to grow cells such as neuronal and primary cells, the protein input needs to be kept rather low. In turn, this also increases the demand on the LC-MS/MS sensitivity. Recently, we have established a chemical proteomics workflow called SP2E that uses carboxylate-modified paramagnetic beads for a protein clean-up and streptavidin-coated paramagnetic beads for an affinity enrichment of the probe-labelled proteins (Figure 1A).^11^ So far, the workflow has been successfully applied for example for identification of ibrutinib-derived probe targeting Bruton’s tyrosine kinase (BTK),^12^ development of vinyl phosphonamidates for cysteine-based protein functionalization^13,14^ and elucidation of Cu(I)-catalyzed azide-alkyne cycloaddition (CuAAC) side-reactivity.^15^

**Figure 1.**
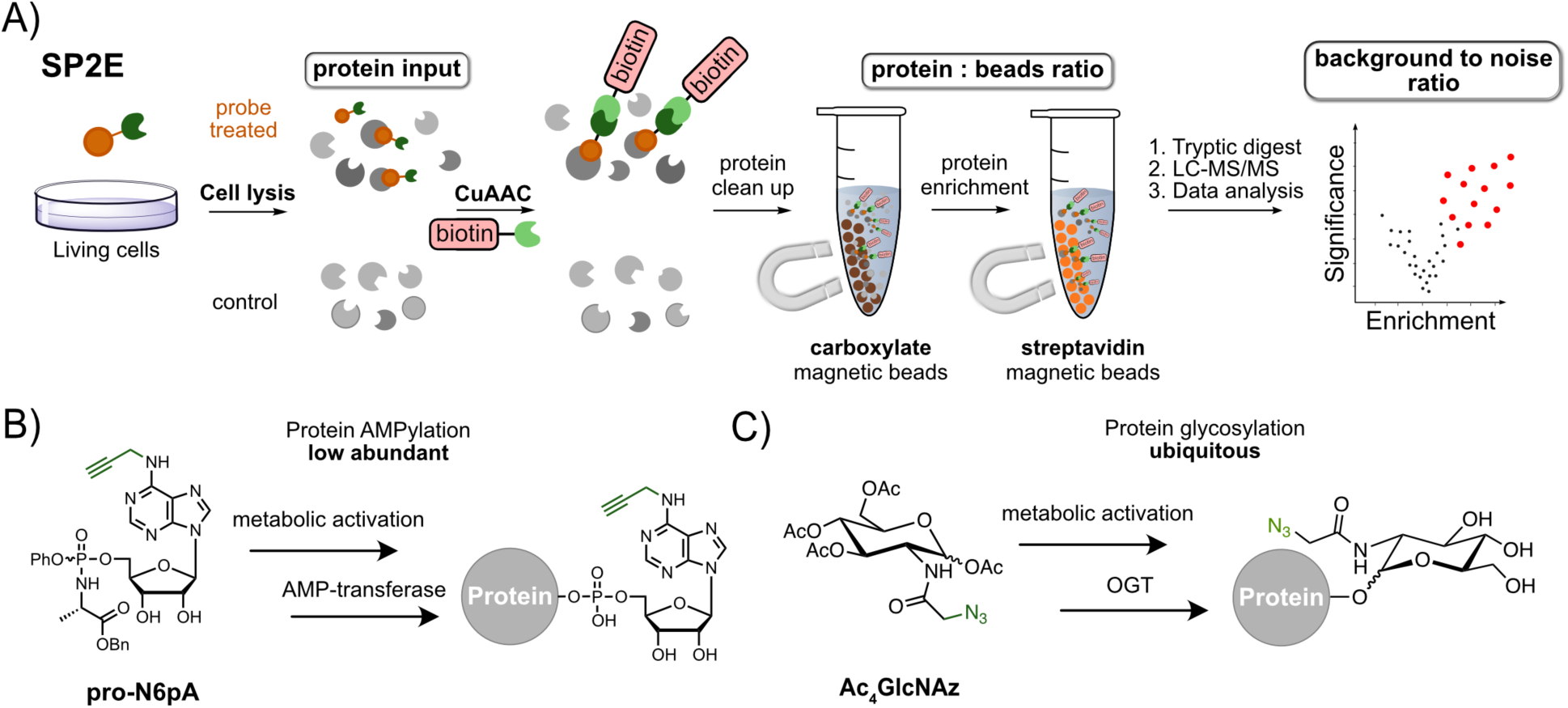
Scaling down and optimization of chemical proteomics SP2E workflow in this study (A). SP2E chemical proteomics workflow. (B) Metabolic labeling of AMPylated and *O*-GlcNAcylated proteins in living cells (C); OGT – *O*-GlcNAc transferase, AMP – adenosine monophosphate.

Here, we describe a development of a small-scale chemical proteomics SP2E workflow to profile low abundant protein PTM called AMPylation (Figure 1B) and ubiquitous protein *O*-GlcNAcylation (Figure 1C) after metabolic labelling from low protein inputs (25 μg). To achieve this, systematic testing of magnetic bead amounts and protein inputs was performed, resulting in significant reduction of magnetic beads consumption and improvement of background-to-noise ratio. The data-independent acquisition (DIA)^16,17^ has proved to be crucial to reach necessary protein coverage and quantification in direct comparison with data-dependent acquisition (DDA) experiments. The workflow is now suitable for a high-throughput low protein input profiling of compound libraries with a minimal hands-on time during sample preparation and straightforward translation to processing by widely available liquid-handling robots.

## MATERIALS AND METHODS

### Cell culture

HeLa (RRID: CVCL_0030), SH-SY5Y (RRID: CVCL_0019) cells were cultured in Dulbecco’s modified Eagle’s medium (DMEM) -- high glucose supplemented with 10% fetal calf serum (FCS) and 2 mM l-alanyl-l-glutamine at 37 °C and a 5% CO_2_ atmosphere.

### Cell seeding and probe treatment

When the confluency of the cells was around 80%, media was removed and cells were washed twice with 4 mL sterile 1x PBS. Afterwards, 1 mL TrypLE^TM^ Express was added and the cells were incubated in CO_2_ incubator for 3 min. The digestion was stopped by adding 4 mL pre-warmed DMEM media. Pipette the cells up and down to get single and evenly distributed cells. The concentration of the cell suspension was determined by mixing 10 µL cell suspension with 10 µL trypan blue using Countess 3 (Invitrogen). Meanwhile, the cell suspension was transferred to a 15 mL falcon and was spun down 3 min at 300 rpm. Afterwards, the supernatant was removed and the cell pellet was resuspended in the DMEM media to the target concentration. 1x 10^6^ SH-S5Y5 cells or 0.6x 10^6^ Hela cells were seeded into each p60 dish, which were incubated in the CO_2_ incubator for 24 h. Then 3 µL Ac_4_GlcNAz (200 mM in DMSO) or pro-N6pA (100 mM in DMSO) was directly added into the media for experimental groups, with 3 µL DMSO added as a control.

### Cell harvest and BCA measurement

Cells were incubated for 16 hours at 37 °C prior to harvesting. Cellular morphology was assessed under an optical microscope before harvest. Cells were then washed twice with 2 mL of Dulbecco’s phosphate-buffered saline (DPBS), scraped into 1 mL of DPBS, and pelleted by centrifugation at 1,000 rpm for 5 minutes at 4 °C. The supernatant was removed, and 200 µL of lysis buffer (20 mM HEPES, pH 7.5, 1% (v/v) NP-40, 0.2% (w/v) SDS) was applied, followed by sonication (10s, at a 20% intensity with a rod sonicator) to initiate cell lysis. The lysate proteins were collected from the supernatant following centrifugation at 13,000 rpm for 10 minutes at 4 °C. Pierce™ BCA Protein Assay Kit was employed for the subsequent measurement of protein concentration. First, a 2 mg/mL bovine serum albumin (BSA) standard was serially diluted to final concentrations of 12.5, 25, 50, 100, 200, and 400 μg/mL, with all samples diluted 40 times to a total volume of 200 µL with MQ water. Triplicates of each 50 µL standard and sample were aliquoted into a clear, flat-bottom 96-well plate, followed by the addition of 100 µL of working reagent (R2:R1 = 1:50). The plate was incubated at 60 °C for 15 minutes, then allowed to cool to room temperature. Absorbance at 562 nm was measured using a Tecan plate reader, and protein concentrations were calculated accordingly.

### Protein enrichment and Digestion

Optimization of the SP2E enrichment protocol was conducted in duplicate while the verification experiment was performed with four biological replicates.

### Click reaction Condition A

Varying amounts of protein, with or without probe treatment, were diluted in lysis buffer (20 mM HEPES, pH 7.5, 1% (v/v) NP-40, 0.2% (w/v) SDS) to a final reaction volume of 40 µL. The master mix containing 0.4 *μ*L of biotin-PEG-N_3_ (10 mM in DMSO), 1.2 *μ*L of TCEP (100 mM in water), 0.25 *μ*L of TBTA (16.7 mM in DMSO) and 3.15*μ*L of MS-grade H_2_O and was added to each sample. Samples were gently vortexed, and the click reaction was initiated by the addition of 5 *μ*L of CuSO_4_ (10 mM in water) and incubated for 1.5 h (25 °C, 450 rpm).

### Condition B

Various amounts of lysates were diluted in lysis buffer (20 mM HEPES, pH 7.5, 1% (v/v) NP- 40, 0.2% (w/v) SDS) to a final reaction volume of 40 µL. A 5 *μ*L master mix containing 0.5 *μ*L of biotin-PEG-N_3_ (10 mM in DMSO, for MS measurement) or 5/6-TAMRA-Azide-Biotin (10 mM in DMSO, for in-gel fluorescence imaging), 1.5 *μ*L of TCEP (200 mM in water), 0.3 *μ*L of TBTA (16.7 mM in DMSO) and 2.7 *μ*L of MS-grade H_2_O and was added to each sample. Samples were gently vortexed, and the click reaction was initiated by the addition of 5 *μ*L of CuSO_4_ (10 mM in water) and incubated for 1.5 h (25 °C, 450 rpm).

### Condition C

400 *μ*g of lysates were diluted in lysis buffer (20 mM HEPES, pH 7.5, 1% (v/v) NP-40, 0.2% (w/v) SDS) to a final reaction volume of 200 µL. A 5.25 *μ*L master mix containing 2 *μ*L of 5/6- TAMRA-Azide-Biotin (10 mM in DMSO), 3 *μ*L of TCEP (200 mM in water), 0.25 *μ*L of TBTA (83.5 mM in DMSO) was added to each sample. Samples were gently vortexed, and the click reaction was initiated by the addition of 2 *μ*L of CuSO_4_ (100 mM in water) and incubated for 1.5 h (25 °C, 450 rpm).

### Condition D

25 *μ*g of lysates were diluted in lysis buffer (20 mM HEPES, pH 7.5, 1% (v/v) NP-40, 0.2% (w/v) SDS) to a final reaction volume of 40 µL. Then, 5 *μ*L of Iodoacetamide (139.5 mM in water, freshly prepared) was added to each sample. The mixture was mixed, spun down, and incubated for 30 min in the dark (25 °C, 750 rpm). Afterwards, the SPAAC click reaction was triggered by the addition of another 5 *μ*L of DBCO-PEG_4_-Biotin (200 *μ*M in water) and incubated for 30 min (25 °C, 750 rpm).

### Protein enrichment

#### Automated workflow (with Hamilton MicroPrep Workstation)

During the click reaction, carboxylate-coated magnetic beads (hydrophobic : hydrophilic = 1:1) and streptavidin-coated magnetic beads were each washed three times with either 500 µL MS-grade water or 500 µL of 0.2% SDS in PBS, respectively. Following completion of the click reaction, 30 µL of 8 M urea was added to each replicate. The reaction mixtures were then transferred to the pre-equilibrated carboxylate-coated magnetic beads in a 96-well PCR plate.

From this point, all subsequent steps were performed using a Hamilton MicroPrep automated workstation. The beads were resuspended, and 100 µL of absolute ethanol was added. The plate was shaken at room temperature for 5 minutes at 950 rpm. After incubation, the supernatant was removed, and the beads were washed three times with 150 µL of 80% ethanol in water, followed by a single wash with 150 µL of LC-MS-grade acetonitrile. Proteins bound to the beads were then eluted three times with 60 µL of 0.2% SDS in PBS at 40 °C for 5 minutes (950 rpm). All eluates were pooled and transferred to pre-washed streptavidin-coated beads, followed by incubation for 1 hour at 25 °C with shaking (950 rpm). The streptavidin beads were then washed three times with 150 µL of 0.1% NP-40 in PBS, twice with 150 µL of freshly prepared 6 M urea, and twice with MS-grade water.

#### Manual workflow

During the click reaction, carboxylate-coated magnetic beads (hydrophobic : hydrophilic = 1:1) and streptavidin-coated magnetic beads were each washed three times with either 500 µL MS-grade water or 500 µL of 0.2% SDS in PBS, respectively. Following completion of the click reaction, 30 µL of 8 M urea was added to each replicate. The reaction mixtures were then transferred to the pre-equilibrated carboxylate-coated magnetic beads in a 96-well PCR plate. The beads were resuspended in ThermoMixer for 1 min (25 °C, 950 rpm), and 100 µL of absolute ethanol was added. The plate was shaken for 5 minutes (25 °C, 950 rpm). After incubation, the supernatant was removed with multichannel pipette, and the beads were washed three times with 150 µL of 80% ethanol in water, followed by a single wash with 150 µL of LC-MS-grade acetonitrile with magnet plate. Proteins bound to the beads were then eluted three times with 60 µL of 0.2% SDS in PBS at 40 °C for 5 minutes (950 rpm). All eluates were pooled and transferred to pre-washed streptavidin-coated beads, followed by incubation for 1 hour at 25 °C with shaking (950 rpm). The streptavidin beads were then washed three times with 150 µL of 0.1% NP-40 in PBS, twice with 150 µL of freshly prepared 6 M urea, and twice with 150 µL MS-grade water.

#### Large scale SP2E workflow

During the reaction, carboxylate-coated magnetic beads (hydrophobic : hydrophilic = 1:1) and streptavidin-coated magnetic beads were each washed three times with either 1 mL MS-grade water or 1 mL of 0.2% SDS in PBS, respectively. Following completion of the click reaction, 200 µL of 8 M urea was added to each replicate. The reaction mixtures were then transferred to the pre-equilibrated carboxylate-coated magnetic beads in 1.5 mL Eppendorf vials, resuspended, and 600 *μ*L of ethanol was added. Subsequently, the mixture was incubated for 5 min (25 °C, 950 rpm), and washed thrice with 500 *μ*L 80% EtOH in water. Between each wash, the mixture was shortly vortexed, spun down and placed on a magnetic rack for 1 min. Thereafter, the protein was eluted from the carboxylate-coated beads to the pre-washed streptavidin-coated beads by incubating 5 min with 500 *μ*L of 0.2% SDS in PBS twice (25 °C, 950 rpm). The combined eluates were incubated for 1 h (25 °C, 950 rpm), followed by thrice wash with 500 *μ*L of 0.1% NP40 in PBS, twice with 500 µL of freshly prepared 6 M urea, and twice with 500 µL MS-grade water.

### On-beads digest of enriched proteins Condition A

The washed beads were resuspended in 50 µL of 50 mM TEAB buffer and digested overnight at 37 °C with 1.5 µL of sequencing-grade trypsin (0.5 mg/mL), with caps securely closed. The following day, the supernatant was transferred to a new microcentrifuge tube. The beads were then washed twice with 20 µL of TEAB buffer at 40 °C for 5 minutes (950 rpm), followed by twice wash with 20 µL of MS-grade water. All wash fractions were combined with the original digest, acidified by the addition of 0.9 µL of formic acid, and transferred directly to MS vials without desalting.

### Condition B

The washed beads were resuspended in 50 µL of 50 mM TEAB buffer and digested overnight at 37 °C with 1.5 µL of sequencing-grade trypsin (0.5 mg/mL), with caps securely closed. The following day, the supernatant was transferred to a new microcentrifuge tube. The beads were then washed once with 20 µL of TEAB buffer at 40 °C for 5 minutes (950 rpm), followed by one wash with 20 µL of MS-grade water. All wash fractions were combined with the original digest, acidified by the addition of 0.9 µL of formic acid, and transferred directly to MS vials without desalting. (For the manual experiment, only once with TEAB buffer and once with MS-grade water)

### In-gel fluorescence analysis

Following the final wash step of the SP2E workflow, 16 µL of MS-grade H_2_O and 4 µL of 5x LB (prepared by freshly mixing 200 µL 5x loading buffer with 20 µL 1M DTT) were added to the streptavidin-coated beads. The mixture was vortexed, spun down, and incubated at 95 °C for 5 minutes. After cooling to room temperature, the supernatants were loaded onto a 10% SDS-PAGE gel. Electrophoresis was performed at 100 V for 10 minutes, followed by 150 V for 80 minutes in 1x running buffer (25 mM Tris, 192 mM glycine, 0.1% SDS in ddH₂O) under dark conditions. Fluorescently labeled proteins were subsequently visualized in-gel using the Amersham ImageQuant™ 800 imaging system (Cytiva).

### Western blotting

After the SDS-PAGE gel running, the separated proteins were transferred onto PVDF membrane using a blotting sandwich moistened by Semi-dry transfer buffer (48 mM Tris, 39 mM glycine, 0.0375% (m/v) SDS, 20% (v/v) methanol). The transfer was carried out 30 min at 25 V (standard method) with TransBlot^®^ Turbo™ Transfer System (Bio-Rad). Afterwards, the PVDF membrane was incubated 60 min in blocking solution (0.5 g milk powder in 10 mL PBST (PBS +0.5% Tween)). Subsequently, 10 *μ*L primary antibody (1:1000 dilution time) with specificity for the protein of interest was added and the mixture was incubated 1 h at 4 °C overnight followed by 3 times wash with PBST for 10 min each time. Then 1 *μ*L of the HRP-conjugated horse-anti mouse IgG(H+L) antibody (1:10000 dilution) and 1 *μ*L hFAB™ Rhodamine Anti-GAPDH Primary Antibody in 10 mL blocking solution were added. After 1 h of incubation at room temperature the membrane was washed again 3 times for 10 min with PBST. Then, 500 *μ*L ECL Substrate and 500 *μ*L peroxide solution (Amersham™ ECL Detection Reagents, Cytiva) were mixed in Eppendorf tube and added to the membrane to stain the Western blot. Finally, images of the Western blot were taken by the Amersham ImageQuant™ 800 imaging system (Cytiva).

### OSMI-1 inhibitor screening experiment

A total of 0.6x 10^6^ Hela cells were seeded into each 60 mm culture dish, and incubated for 24 h in the CO_2_ incubator to allow cell attachment and recovery. Subsequently, the culture medium was replaced with 1.5 mL of fresh medium containing the respective treatment conditions: DMSO control or OSMI-1 at final concentrations of 20 nM, 100 nM, 200 nM, 1000 nM, 2000 nM, or 4000 nM. Each condition was prepared in duplicate and incubated for 3 h. Following the initial OSMI-1 treatment, one dish from each concentration group received an additional 1.5 mL of fresh medium supplemented with Ac_4_GlcNAz to achieve a final concentration of 200 µM. The corresponding paired dish received an equal volume of DMSO-containing medium as a control. All dishes were subsequently incubated for an additional 24 h in the CO_2_ incubator. Two independent biological replicates were performed for each treatment condition, resulting in a total of 28 culture dishes. Cell harvesting and protein concentration determination were carried out as described above. Subsequently, 25 µg of protein was collected twice from each dish to generate two technical replicates per sample. The resulting samples were then processed using the above-described SP2E workflow and analyzed by data-independent acquisition (DIA) mass spectrometry.

### LC-MS/MS measurement – Thermo Scientific™ Orbitrap Eclipse™ Tribrid™ MS

MS measurements were performed on a Thermo Scientific™ Orbitrap Eclipse™ Tribrid™ MS (Thermo Fisher Scientific) coupled to an UltiMate™ 3000 Nano-HPLC (Thermo Fisher Scientific)) via a nanospray Flex ion source (Thermo Fisher Scientific) equipped with column oven (Sonation) and FAIMS interface (Thermo Fisher Scientific). First, peptides were loaded on an Acclaim PepMap 100 μ-precolumn cartridge (5 μm, 100 A, 300 μm ID x 5 mm, Thermo Fisher Scientific), followed by separation at 40 °C on a PicoTip emitter (noncoated, 15 cm, 75 μm ID, 8 μm tip, New Objective) that was in house packed with Reprosil-Pur 120 C18-AQ material (1.9 *μ*m, 150 A, Dr. A. Maisch GmbH). The HPLC gradient was run from 4-35.2% acetonitrile supplemented with 0.1% formic acid (Buffer B) together with MS-grade H2O supplemented with 0.1% FA (Buffer A) during method (0-5min 4%, 5-6 min to 7%, 7-36 min to 24.8%, 37-41 min to 35.2%, 42-46 min 80%, 47-60 min 4%) at a flow rate of 300 nL/min. Injection volume is 5 μL.

### DIA measurement method – Thermo Scientific™ Orbitrap Eclipse™ Tribrid™ MS

FAIMS was performed with one compensation voltage (CV) at -45 V for the whole duty cycle at positive mode. The data-independent acquisition duty cycle consisted of one MS1 scan followed by 30 MS2 scans with an isolation window of 4 m/z range, overlapping with an adjacent window at the 2 m/z range. MS1 scan was conducted with Orbitrap at 60,000 resolution power and a scan range of 200 -1800 m/z and an adjusted RF lens at 30% with standard AGC target and 50ms maximum injection time. MS2 scans were conducted with Orbitrap at 30,000 resolution power, RF lens was set to 30%. The precursor mass window was restricted to a 500 – 740 m/z range. HCD fragmentation was enabled as an activation method with a fixed collision energy of 35%. Normalized AGC target was set to 200% with automatic maximum injection time.

### LC-MS/MS measurement – Thermo Scientific™ Orbitrap™ Astral MS

MS measurements were performed on a Thermo Scientific™ Orbitrap™ Astral MS coupled to an Thermo Scientific™ Vanquish™ Neo™ via an easy spray ion source (Thermo Fisher Scientific) equipped with FAIMS interface (Thermo Fisher Scientific). Peptides were separated at 50 °C with an Ionoptics Aurora column (75 μm ID x 25cm). The 24min HPLC gradient was run with front-loading at 450 nl/min from 1-12% acetonitrile supplemented with 0.1% formic acid (Buffer B) together with MS-grade H_2_O supplemented with 0.1% FA (Buffer A) followed by 17.5 min separation from 12-40% B at 200 nl/min and a 4.5 min wash at 99% B at 300 nl/min. Injection volume varied from 0.1 to 5 uL.

### DIA measurement method – Thermo Scientific™ Orbitrap™ Astral MS

MS1 scan was performed at a resolution of 240K with a scan range from 380-980 m/z. A single compensation voltage of -45V was used for FAIMS. For MS2 the DIA window size and injection time was adjusted according to the load with 10 m/z and 20 ms for higher loads and 20 m/z and 40 ms for the lower load injections.

### DDA measurement method – Thermo Scientific™ Orbitrap Eclipse™ Tribrid™ MS

FAIMS was switched between two CV at -50 V and -70 V for MS1 scan during the 1.7s duty cycle at positive mode. The data-dependent acquisition duty cycle consisted of one MS1 scan followed by as many MS2 scans as possible until the cycle time reached 1.7s. MS1 scan was conducted with Orbitrap at 240,000 resolution power and a scan range of 375 -1500 m/z with an adjusted RF lens at 30%. MS1 AGC target is standard and MS1 maximum injection time is 50ms. MS2 scans were conducted with rapid Ion trap scan rate with RF lens set to 30%. HCD fragmentation was enabled as an activation method with a fixed collision energy of 30%. Standard MS2 AGC target was applied with 35ms as MS2 maximum injection time. For MS2 quadrupole isolation window, 1.2 m/z was set without isolation offset. The intensity threshold is 1.0e4 counts with included charge states 2-6 and dynamic exclusion 40s.

### Data analysis DIA data

Raw files were converted in the first step with “MSConvertGUI” ^18^ as a part of the “ProteoWizard” software package (http://www.proteowizard.org/download.html) to an output mzML format applying the “peakPicking” filter with “vendor msLevel=1”, and the “Demultiplex” filter with parameters “Overlap Only” and “mass error” set to 10 ppm. Standalone DIA-NN software under version 1.8.1 or 1.9 or 2.1.0 (always the latest released version during the data analysis) was used for protein identification and quantification.^5^ First, a spectral library was predicted in silico by the software’s deep learning-based spectra, RTs and IMs prediction using Uniprot Human FASTA (containing canonical and isoforms, decoys, common contaminants). DIA-NN search settings: FASTA digest for library-free search/library generation option was enabled, together with a match between runs (MBR) option (not available in DIA-NN version 1.8.1 and 1.9) and precursor FDR level set at 1%. Library generation was set to smart profiling, Quantification strategy - Robust LC. The mass accuracy and the scan window were set to 0 to allow the software to identify optimal conditions. The precursor m/z range was changed to 500-740 m/z to fit the measuring parameters.

Perseus (1.6.10.43) was used to log_2_ transform LFQ intensities, and then sample rows were assigned to two groups including either control or probe treated. Next, the annotated rows were filtered for at least 2 valid values out of duplicates experiments or at least three valid values out of four replicates experiments in at least one group. Besides, missing values were replaced from normal distribution. -log10(p-values) were obtained by a two-sided one sample Student’s t-test over replicates with the initial significance level of p = 0.05 adjustment by the multiple testing correction method of Benjamini and Hochberg (FDR = 0.05).

### DDA data

Raw files were converted with “FAIMS MzXML Generator”, which split a FAIMS Raw file into a set of MaxQuant compliant MzXML files, each containing only scans collected using a single correction voltage, into MzXML files. MaxQuant software under version 2.7 was used for protein identification and quantification with Andromeda search engine.^19^ “Standard” was chosen as analysis type and Oxidation and Acetyl (Protein N-term) ware used as variable modifications with “Max. number of modifications per peptide” as 5. Searches were performed against the Uniprot database for Homo sapiens (taxon identifier: 9606, 5th November 2025, including isoforms). At least one unique peptide was required for protein identification. Protein quantification was conducted with MaxLFQ method with a minimum ratio count of two using both unique and razor peptides. False discovery rate determination was carried out using a decoy database and thresholds were set to 1% FDR both at peptide-spectrum match and at protein levels.

### Data Visualization

Volcano plot was performed using Perseus.1.6.10.43.^20^ For statistical analysis, the protein matrix was filtered to retain proteins with at least three valid values in at least one group (Ac_4_GlcNAz-treated group or DMSO-treated group). Missing values were imputed from a normal distribution using a width of 0.3 and a downshift of 1.8. Heatmap analysis was generated using the “Difference” parameter as the color scale. Proteins identified as significantly upregulated in the DMSO-Glc versus DMSO-DMSO volcano plot were used as the x-axis entries, while the different treatment conditions (group means) were displayed on the y-axis. For this analysis, the log_2_-transformed LFQ intensities of the significantly upregulated proteins were extracted from six additional OSMI-1 treatment conditions. In the middle of data extraction, missing values were imputed using a custom approach. Specifically, if a protein intensity value was missing, it was replaced with the globally minimum detected log_2_-(LFQ intensity) value within the corresponding dataset, followed by subtraction of 1.5 to simulate low-abundance protein signals below the detection threshold. For mean-based comparisons, the difference values were calculated by subtracting the mean log_2_- (LFQ intensity) of the control group from the mean log_2_-(LFQ intensity) of the corresponding Ac_4_GlcNAz-treated group.

## RESULTS AND DISCUSSION

To enable low protein input chemical proteomics applications, we aimed to scale down the published SP2E workflow (Figure 1A) while maintaining robust protein recovery and enrichment performance. This posed a substantial challenge, as workflow miniaturization required systematic evaluation of series of experimental parameters, including different combinations of carboxylate magnetic beads for protein clean-up and streptavidin-coated beads for affinity enrichment with a decreasing protein input. The large number of possible bead pairings, each with distinct binding capacities, necessitated iterative optimization to identify conditions compatible with reduced sample input without compromising reproducibility or proteome coverage. To evaluate the performance of the scaled-down workflow, we selected two distinct protein PTMs that can be addressed using established chemical proteomic probes. As a low-abundance target, we focused on protein AMPylation using the probe pro-N6pA, a PTM that is typically difficult to detect due to its low stoichiometry (Figure 1B). In parallel, we investigated the more abundant protein *O*-GlcNAcylation using the metabolic labeling by Ac_4_GlcNAz (Figure 1C). Together, these complementary systems enabled assessment of the workflow across markedly different PTM abundance levels and enrichment demands. The optimization of the method was carried out using the liquid handling robot from Hamilton (MicrolabPrep). We first focused on optimizing protein AMPylation analysis in HeLa cells by titrating the protein input from 25 to 100 µg while initially maintaining the bead quantities reported in the original SP2E protocol, namely 5000 µg of carboxylate magnetic beads for protein clean-up and 200 µg of streptavidin-coated beads for affinity enrichment. All measurements were performed using a DIA mass spectrometry method. Analysis of the four marker proteins ABHD6 (α/β hydrolase domain-containing protein 6), ACP2 (lysosomal acid phosphatase 2), PLD3 (phospholipase D3), and PPME1 (protein phosphatase methylesterase 1) revealed markedly different enrichment efficiencies under these conditions. While enrichment of ABHD6 and ACP2 improved upon decreasing the protein input, recovery of PPME1 and PLD3 remained comparatively poor, indicating protein-dependent differences in enrichment performance (Figure 2A). Next, we systematically reduced the amounts of both carboxylate magnetic beads and streptavidin-coated beads while maintaining a constant protein input of 100 µg. Notably, decreasing the streptavidin bead quantities from 200 to 80 µg had no discernible impact on the overall enrichment efficiency (Figure 2B). However, further decrease of the amount of streptavidin beads lead to either no enrichment or signal loss of the marker proteins suggesting unfavorable signal-to-noise ratio and insufficient biding capacity (Figure S1). In contrast, decreasing the amount of carboxylate beads led to a pronounced reduction in background signals for PPME1, indicating an improvement in enrichment specificity under these conditions (Figure 2C).

**Figure 2.**
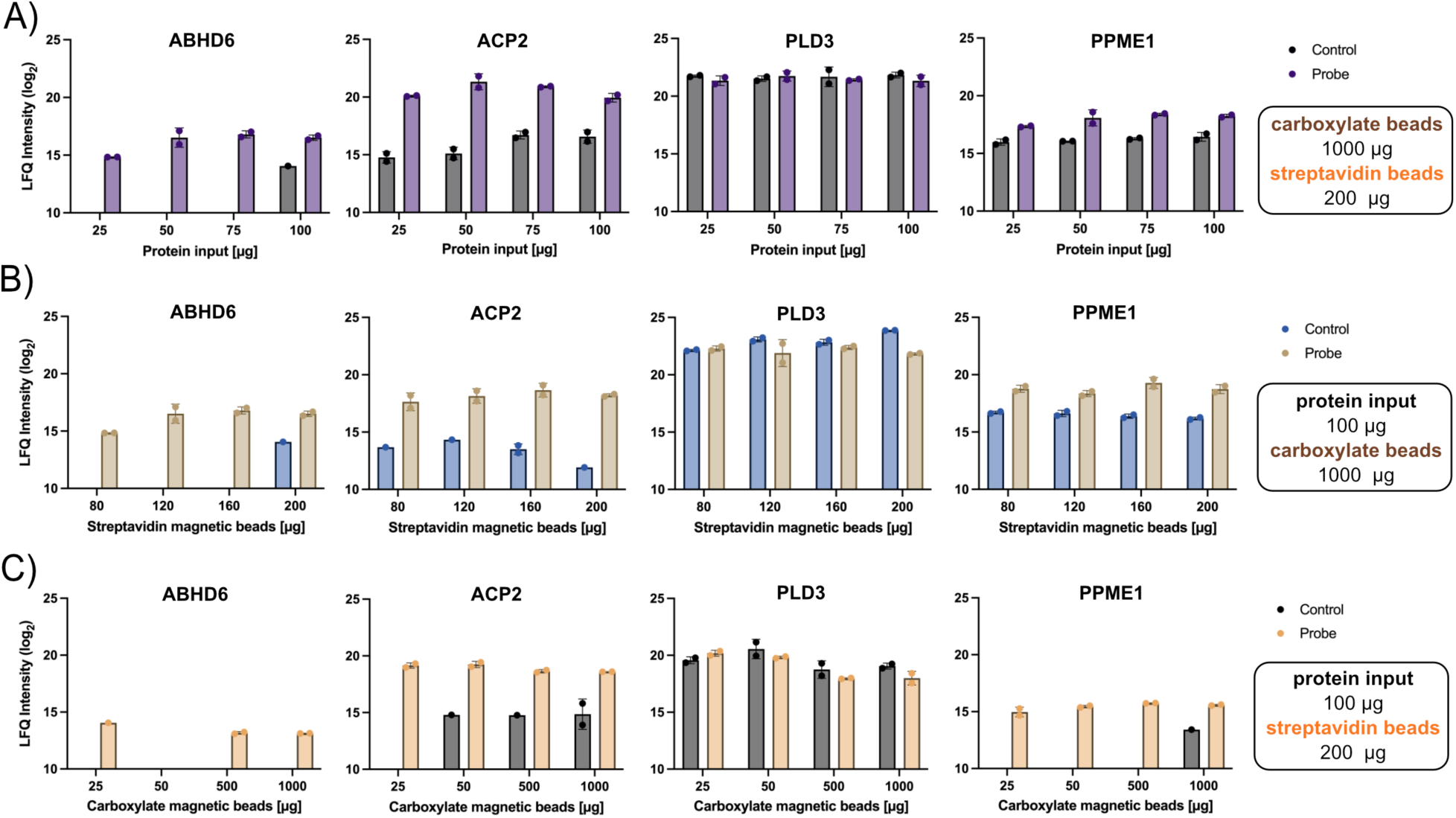
Iterative scaling down of input protein amounts (A) combined with decreased loading of carboxylate- (B) and streptavidin-coated (C) magnetic beads. All treatments with pro-N6pA probe were done on HeLa cells in two biological replicates.

Next, we tested if click reaction with a faster kinetics would lead to a better outcome. Therefore, we used picolyl-azide-biotin reagent instead of more broadly used alkyl-azide-based reagent.^21^ However, picolyl-azide-biotin has resulted in the same enrichment efficiency as standard alkyl-azide (Figure 3A). While the MS/MS measurements were done on Orbitrap™ Eclipse using a DIA method employing Orbitrap for both MS1 and MS2, we next tested how the instrument sensitivity and speed would influence the outcome of the SP2E workflow. The same enriched sample set from control and probe treated cells (100 µg input protein, 25 µg carboxylate and 80 µg streptavidin beads) was measured in parallel on Orbitrap Eclipse and Orbitrap™ Astral (Figure 3A and B).^6,22^ For the measurement on Astral, the original MS samples were 5- to 50-times diluted. The two instruments produced qualitatively comparable results overall, with all marker proteins identified in every measurement. Despite this, the Astral showed substantially higher sensitivity, identifying approximately twice as many peptides at 5-times dilution and significantly more at 50-times dilution than the Orbitrap Eclipse (Figure 3B). The measurement on Astral was done using a 24min LC-gradient, while on Eclipse a 60min gradient was used. That demonstrates the possibility to significantly increase in the throughput of the chemoproteomics measurements.

**Figure 3.**
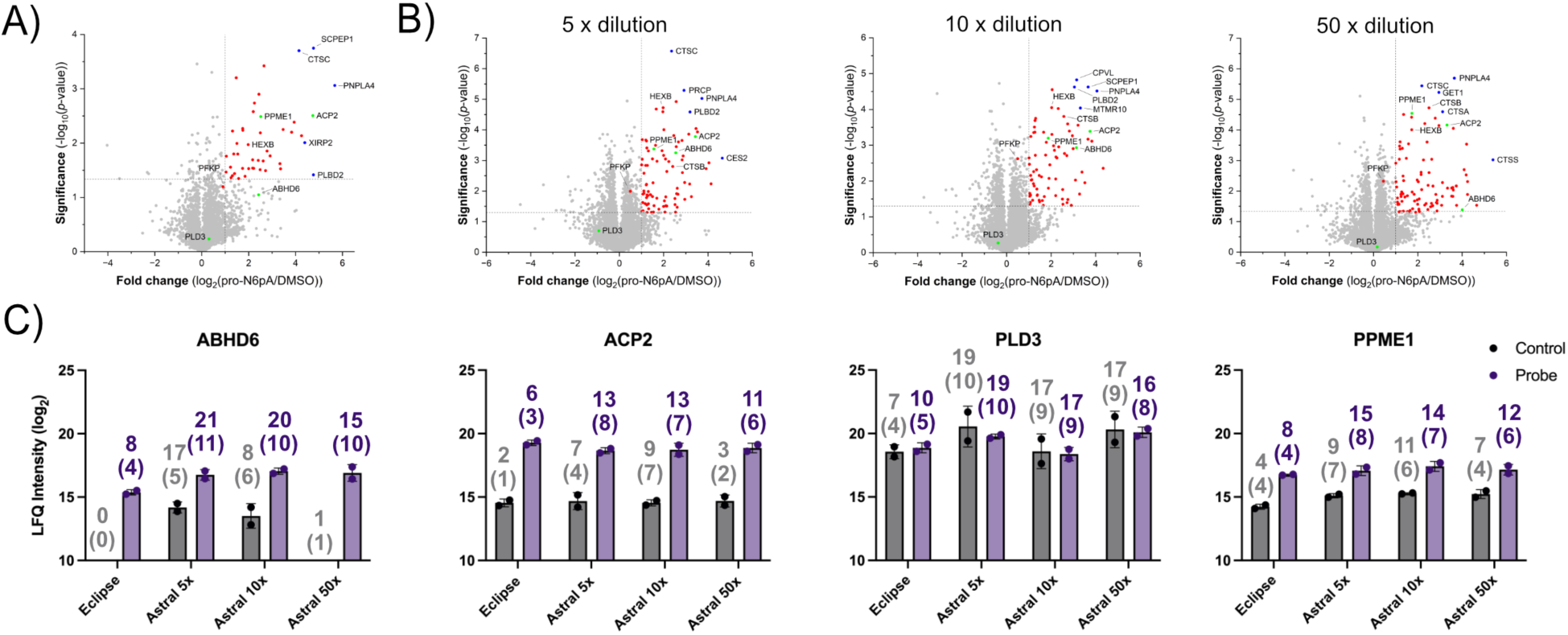
Comparison of MS/MS acquisition on Orbitrap Eclipse and Astral. (A-B) Volcano plots visualizing the enrichment of pro-N6pA labelled proteins (red circles) from HeLa cells using the SP2E workflow starting with 100 µg input protein, utilizing 25 µg carboxylate and 80 µg streptavidin beads, and measured on Orbitrap Eclipse (A) or Astral (B). The four benchmark proteins are shown as green circles (*n* = 4, cut-off p-value < 0.05, and fold-change > 1). (C) Bar plots showing the LFQ intensities from control and pro-N6pA treated HeLa cells measured on Orbitrap Eclipse or Astral with indication of total number of identified peptides and unique peptides (in brackets). All treatments with pro-N6pA probe were done on HeLa cells in two biological replicates.

Because variations in streptavidin beads input can be lowered for still efficient enrichment of the marker proteins, while reduced amounts of carboxylate magnetic beads improved enrichment by lowering background signals, we next selected the lowest working bead amounts (25 µg of carboxylate beads and 80 µg of streptavidin beads) for further optimization. Using these conditions, we again titrated the protein input from 25 to 100 µg, which resulted in satisfactory enrichment of all four benchmark proteins across the tested range (Figure 4A). Subsequently, we applied these optimized conditions to analyze in parallel AMPylated proteins isolated from pro-N6pA–treated SH-SY5Y neuroblastoma and HeLa cells to validate robustness in an independent cellular system (Figure 4B and Figure S2). This confirmed efficient enrichment of all marker proteins, with strong reproducibility across four biological replicates (Figure 4B and 4C). Detailed analysis of the number of identified peptides per protein, including unique peptide counts, further supported the high enrichment efficiency of the optimized workflow (Figure 4D). Comparison of AMPylated proteins found in HeLa and SH-SY5Y cells revealed 39 shared proteins (Figure S3). To additionally demonstrate the sensitivity of the LC–MS/MS-based enrichment approach, we compared the overall scale of pro-N6pA–labelled proteins using in-gel fluorescence analysis, where a clearly detectable signal was only observed at a 400 µg protein input (Figure 4E and Figure S4). Together, these results demonstrate the efficiency and robustness of the optimized SP2E workflow.

**Figure 4.**
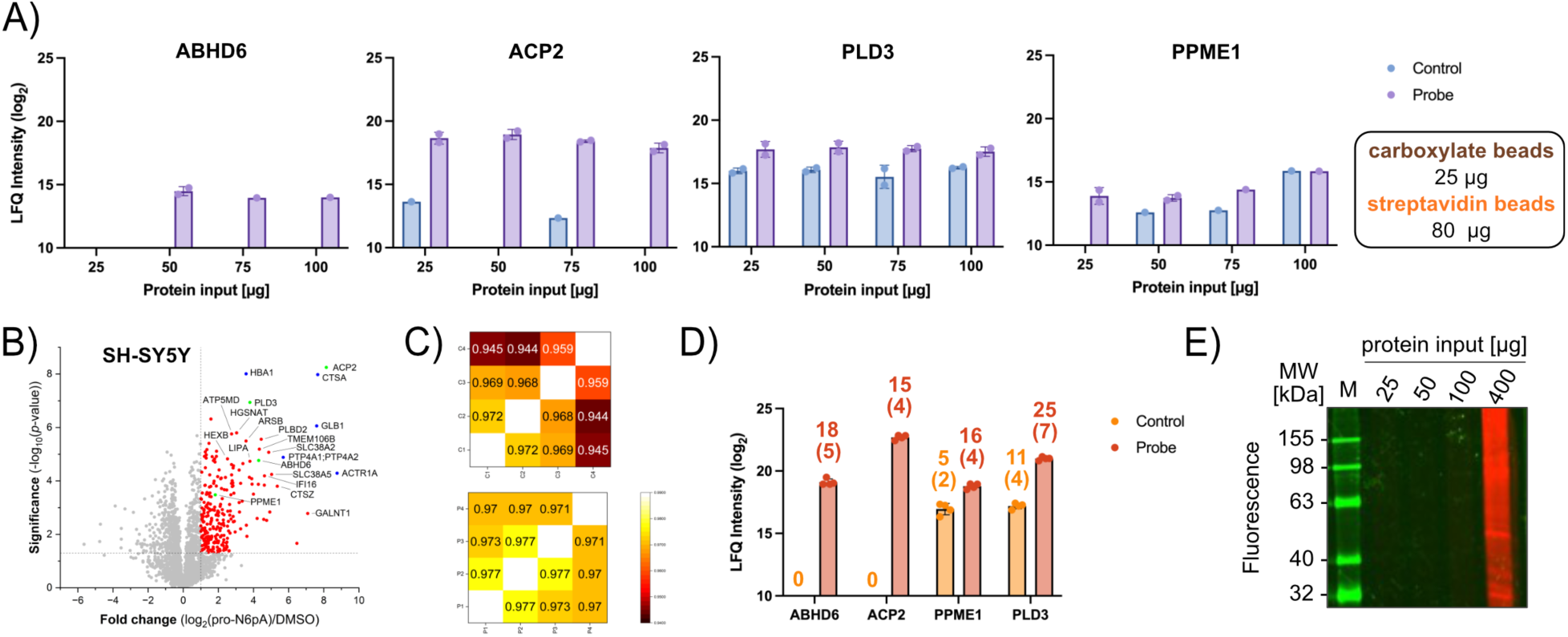
Profiling of AMPylated proteins in living cells. (A) Scaling down the protein input from HeLa cells treated with pro-N6pA with optimized amounts of carboxylate and streptavidin magnetic beads. (B) Volcano plot visualizing the enrichment of pro-N6pA labelled proteins (red circles) from SH-SY5Y cells. The four benchmark proteins are shown as green circles (*n* = 4, cut-off p-value < 0.05, and fold-change > 1). C) Heat maps showing the correlations of label free quantification (LFQ) intensities between the replicates. D) Bar plot showing the LFQ intensities from control and pro-N6pA treated SH-SY5Y cells with indication of total number of identified peptides and unique peptides (in brackets). E) Sensitivity of the in-gel fluorescence analysis for comparison with MS-based analysis in SH-SY5Y cells.

In parallel to the analysis of AMPylated proteins in HeLa cells we extended the optimized workflow to the profiling of *O*-GlcNAcylated proteins, first in HeLa cells and subsequently in SH-SY5Y cells (Figure S5 and Figure 5A). Using a 25 µg protein input, this approach resulted in the significant enrichment of a total of 423 proteins in HeLa and 909 proteins in SH-SY5Y cells, with strong reproducibility and good correlation across biological replicates (Figure 5B). Further inspection of peptide-level identifications, including the canonical *O*-GlcNAc marker protein nuclear pore glycoprotein p62 (NUP62) as well as the five most significantly enriched glycosylated proteins including myomegalin (PDE4DIP), telomere-associated protein RIF1 (RIF1), poly(A)-specific ribonuclease PARN (PARN), lymphokine-activated killer T-cell-originated protein kinase (PBK) and insulin receptor substrate 2 (IRS2) demonstrated excellent sequence coverage and minimal to no detectable background (Figure 5C), underscoring the specificity and robustness of the optimized SP2E workflow.

**Figure 5.**
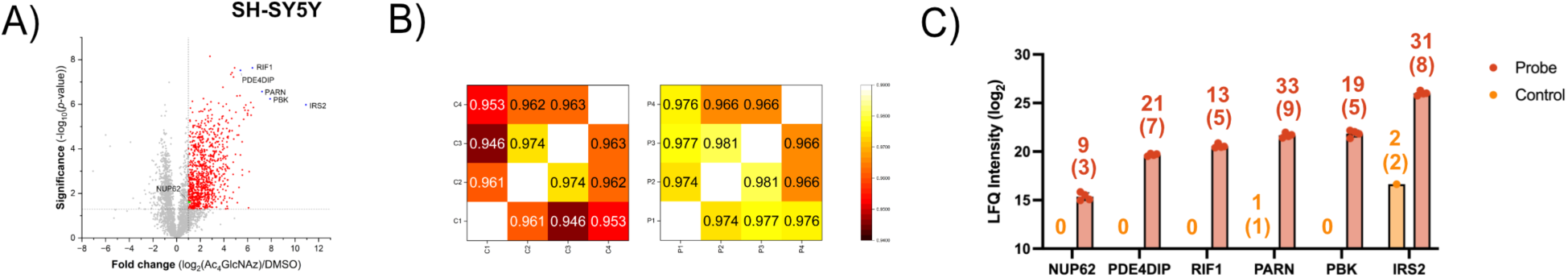
Profiling of *O*-GlcNAcylated proteins in living cells. A) Volcano plot visualizing the enrichment of *O*-GlcNAcylated proteins (red circles) from SH-SY5Y cells (*n* = 4, cut-off p-value < 0.05, and fold-change > 1). The benchmark protein NUP62 (green circle) and five most significantly enriched proteins (blue circles) are marked. B) Heatmap illustrating Pearson correlations among samples within each group. C) Bar plot showing the LFQ intensities from control and Ac_4_GlcNAz treated SH-SY5Y cells with indication of total number of identified peptides and unique peptides (in brackets).

Furthermore, we directly compared DIA and DDA mass spectrometry for both AMPylated and *O*-GlcNAcylated proteome analyses. Overall, DIA resulted in an approximately four-fold increase in the total number of identified proteins (Figure 6A). However, a substantially larger difference was observed at the level of specifically enriched proteins (Figure 6B). In the case of AMPylated proteins, DDA identified only two significantly enriched targets, whereas DIA enabled the detection of approximately 78 enriched AMPylated proteins, corresponding to a 39-fold increase (Figure 6B). For *O*-GlcNAcylated proteins, a similarly strong improvement was observed, with an approximately 14-fold increase in enriched protein identifications with DIA measurements (Figure 6B). Importantly, the number of background proteins remained comparable between both analyses (AMPylation and *O*-GlcNAcylation), indicating that the increased coverage in DIA primarily reflects improved detection of true enriched species rather than increased nonspecific background (Figure 6A). Together, these results highlight the higher sensitivity and fidelity of the DIA-based approach for chemical proteomics applications.

**Figure 6.**
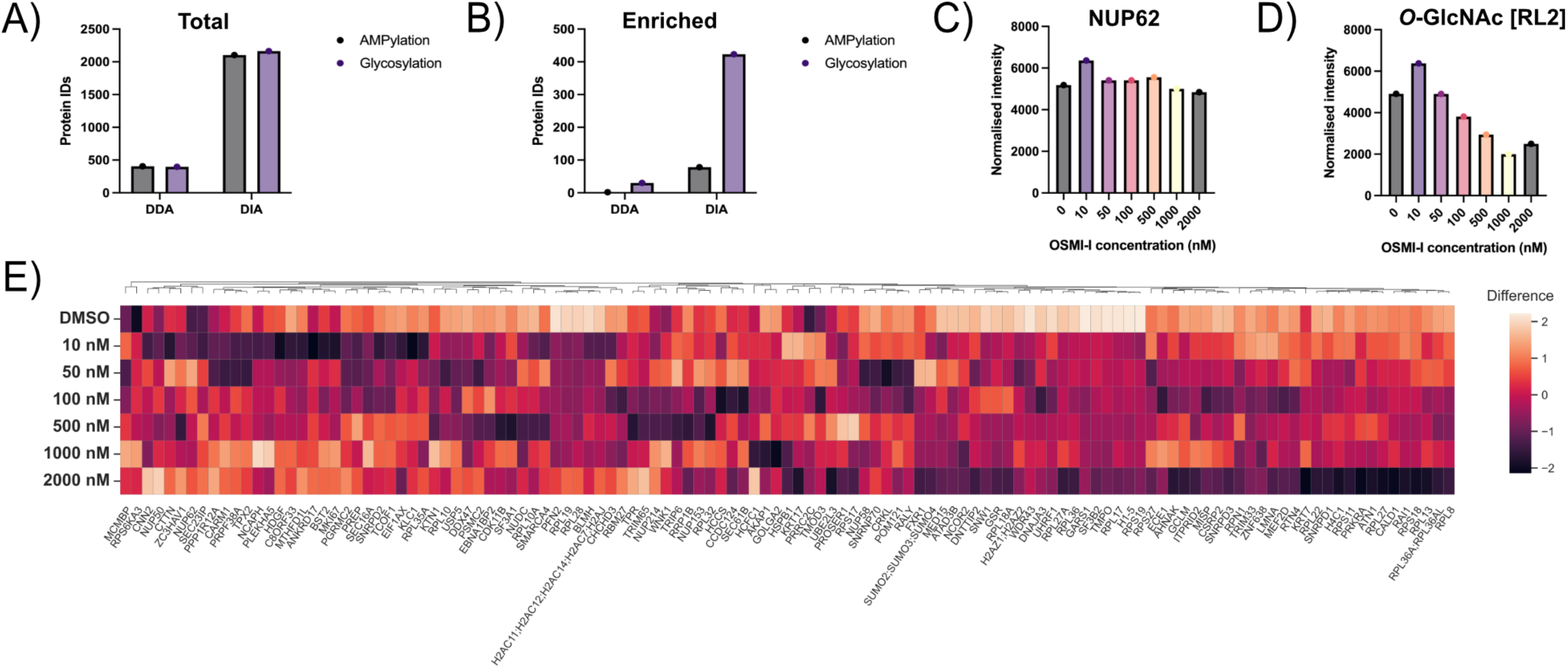
Comparison of DIA and DDA measurements influence on total (A) and enriched (B) protein identifications (IDs). C) Quantification of glycosylated NUP62 from Western blot. D) Overall levels *O*-GlcNAcylated proteins determined by Western blot corroborating the SP2E LC-MS/MS based results. E) Heatmap showing the changes in protein *O*-GlcNAcylation extent after the treatment with different concentrations of OSMI-I, *n* = 4.

Finally, we applied the miniaturized SP2E workflow to profile *O*-GlcNAcylated proteins following inhibition of *O*-GlcNAc transferase (OGT) using OSMI-1 across five different concentrations (Figure S6). In this experimental setup, NUP62 levels remained largely unaffected by OSMI-1 treatment, a finding that was independently confirmed by Western blot analysis (Figure 6C and Figure S7). In contrast, an overall decrease in global *O*-GlcNAcylated protein levels was observed (Figure 6D), as also detected by immunoblotting using the RL2 antibody (Figure S8), confirming effective pathway inhibition under the applied conditions, which was also observed from the heatmap visualizing the decrease of protein *O*-GlcNAcylation (Figure 6E). Together, the OSMI-I treatment experiment suggests only moderate decrease of overall protein *O*-GlcNAcylation that can be monitored by Ac_4_GlcNAz probe.

## CONCLUSIONS

We established a robust small-scale SP2E workflow for chemical proteomics-based profiling of protein post-translational modifications, enabling sensitive and reproducible analysis with substantially reduced protein input. Systematic optimization of key parameters demonstrated that reducing carboxylate magnetic bead amounts by 40-fold improves enrichment efficiency, while streptavidin bead input can be reduced by 2.5-fold. The acquisition of the spectra by Astral provides a better peptide and protein coverage and opens the possibility for further downscaling of the protocol. Under optimized conditions, the workflow enabled reliable detection of both low-abundance AMPylation and abundant *O*-GlcNAcylation across HeLa and SH-SY5Y cells, with strong peptide coverage and high inter-replicate consistency. Benchmarking of mass spectrometry acquisition strategies further highlighted the clear advantage of DIA over DDA, resulting in substantially increased identification of specifically enriched proteins without a corresponding increase in background signals, thereby improving both sensitivity and fidelity of PTM detection. Overall, these findings place the optimized workflow within the broader context of chemical proteomics for post-translational modification analysis, demonstrating that careful downscaling combined with optimized enrichment and DIA-based acquisition provides a scalable and robust platform for PTM profiling from limited biological material.

## ASSOCIATED CONTENT

### Supporting Information

The following files are available free of charge.

Supporting Information (PDF) contains supporting figures and complete list of reagents/materials used in the study.

## Author Contributions

P.K. conceived the study and analyzed the data. L.Z. and J.G. designed and performed the experimental work, mass spectrometric measurements on Orbitrap Eclipse and contributed to data analysis. J. R. set up and carried out measurements on Orbitrap™ Astral. The manuscript was written through contributions of all authors. All authors have given approval to the final version of the manuscript. L.Z. and J.G. designed and performed the experimental work, mass spectrometric measurements and contributed to data analysis.

## Funding Sources

This work was supported by the Deutsche Forschungsgemeinschaft (DFG, German Research Foundation) SFB1309 – 325871075, Boehringer Ingelheim Foundation – Plus 3 Program to P.K. and CSC Scholarship to L.Z.

## Notes

The authors declare no competing financial interest.

## ACKNOWLEDGMENT

We would like to thank our colleagues at the Institute of Chemical Epigenetics, and within the Collaborative Research Center 1309 (DFG) for outstanding institutional support of the project and organization of the measurement time.

## ABBREVIATIONS

ABHD6: α/β hydrolase domain-containing protein 6
ACP2: lysosomal acid phosphatase 2
AMP: adenosine monophosphate
BTK: Burton’s tyrosine kinase
CuAAC: Cu(I)-catalyzed azide-alkyne cycloaddition
DDA: data-dependent acquisition
DIA: data-independent acquisition
IRS2: insulin receptor substrate 2
LC: liquid chromatography
LFQ: label free quantification
MS: mass spectrometry
NUP62: protein nuclear pore glycoprotein p62
OGT: O-GlcNAc transferase
PARN: poly(A)-specific ribonuclease PARN
PBK: lymphokine-activated killer T-cell-originated protein kinase
PDE4DIP: myomegalin
PLD3: phospholipase D3
PPME1: protein phosphatase methylesterase 1
PTM: post-translational modification
RIF1: telomere-associated protein RIF1
TMT: tandem mass tags.

## Notes

### Competing Interest Statement

The authors have declared no competing interest.

