## Supporting Information for "A Scalable and Robust Workflow for Cost-Effective Post-Translational Modifications Profiling by Chemical Proteomics"

**SUPPORTING FIGURES**

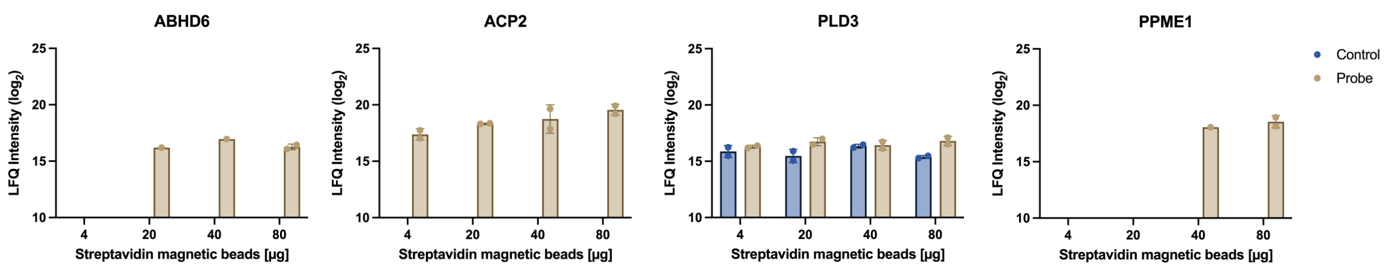

**Figure S1.** Scaling down amount of streptavidin-coated magnetic beads for enrichment of AMPylated proteins in HeLa cells.

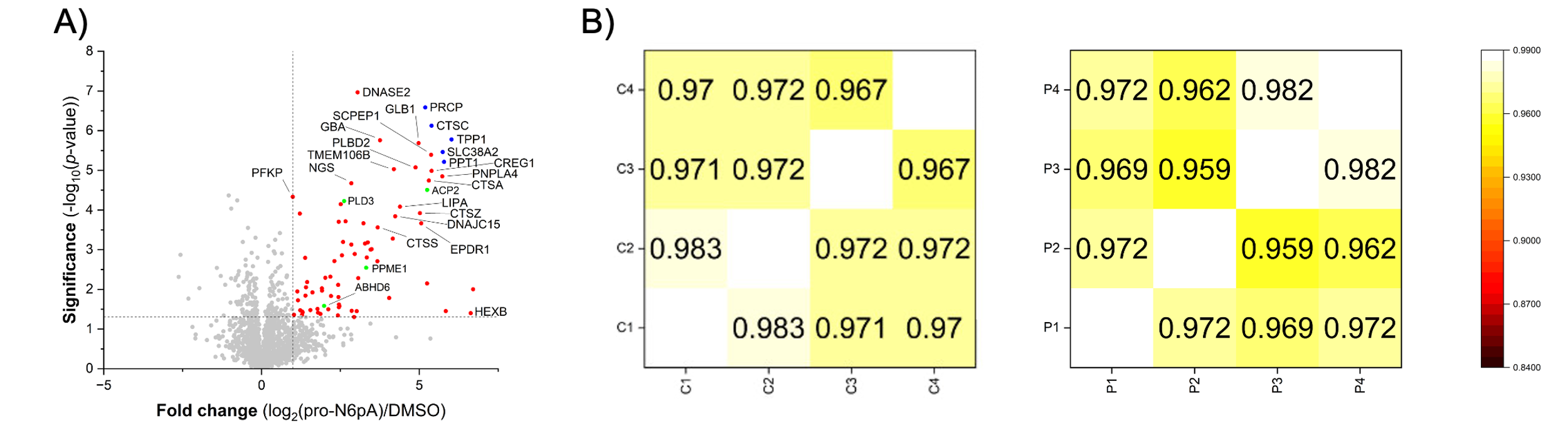

**Figure S2.** Volcano plot visualizing the enrichment of pro-N6pA labelled proteins (red circles) from Hela cells. A) The four benchmark proteins are shown as green circles and the top five significantly enriched proteins are marked as blue (n = 4, cut-off p-value < 0.05, and fold-change > 1). B) Heatmap illustrating Pearson correlations among samples within each group.

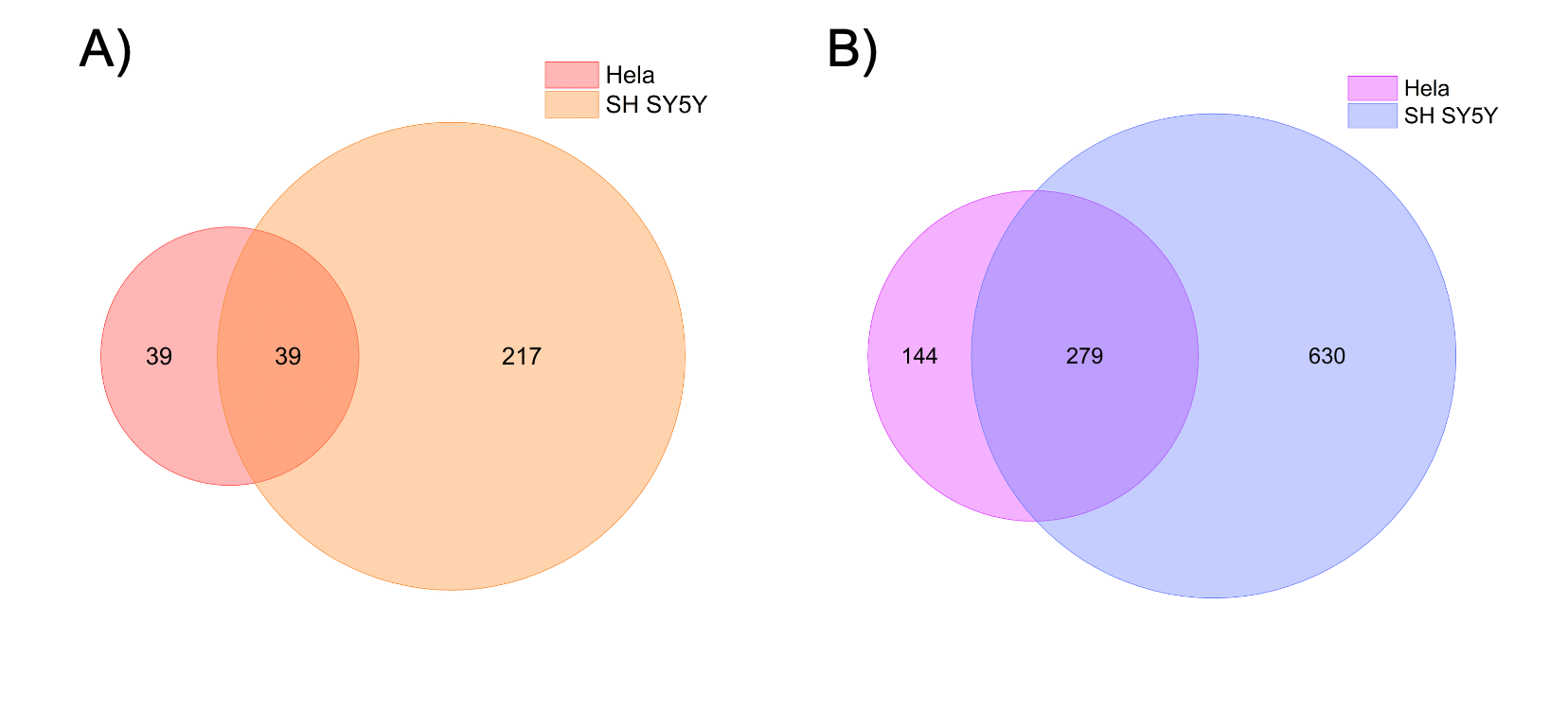

**Figure S3.** Venn diagram illustrating the overlap of all significantly enriched modified proteins identified in HeLa and SH-SY5Y cells using a 25 µg protein input. A) Significantly enriched proteins identified using the pro-N6pA probe. B) Significantly enriched *O*-GlcNAcylated proteins identified using the Ac_4_GlcNAz probe.

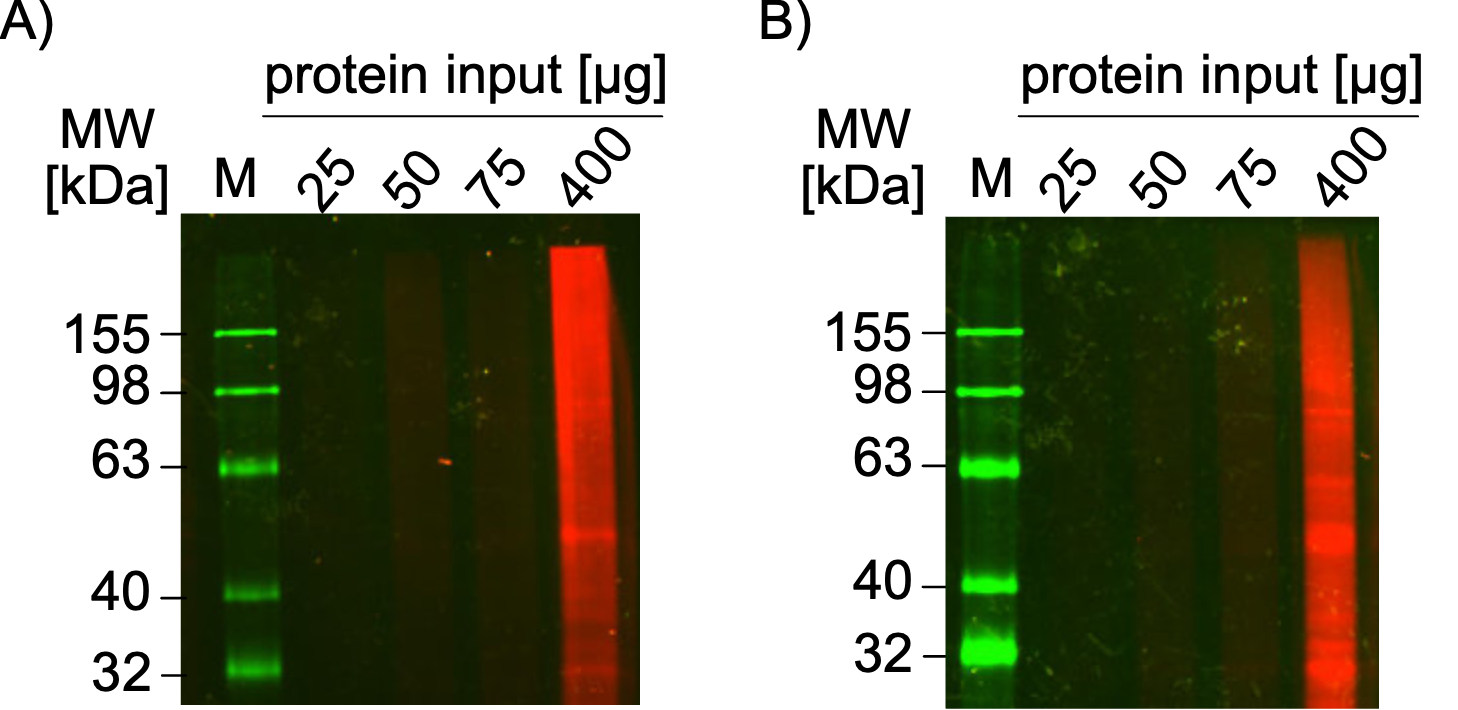

**Figure S4.** Sensitivity of the in-gel fluorescence analysis for comparison with MS-based analysis in HEK293T cells (A) and Hela cells (B) with different protein input.

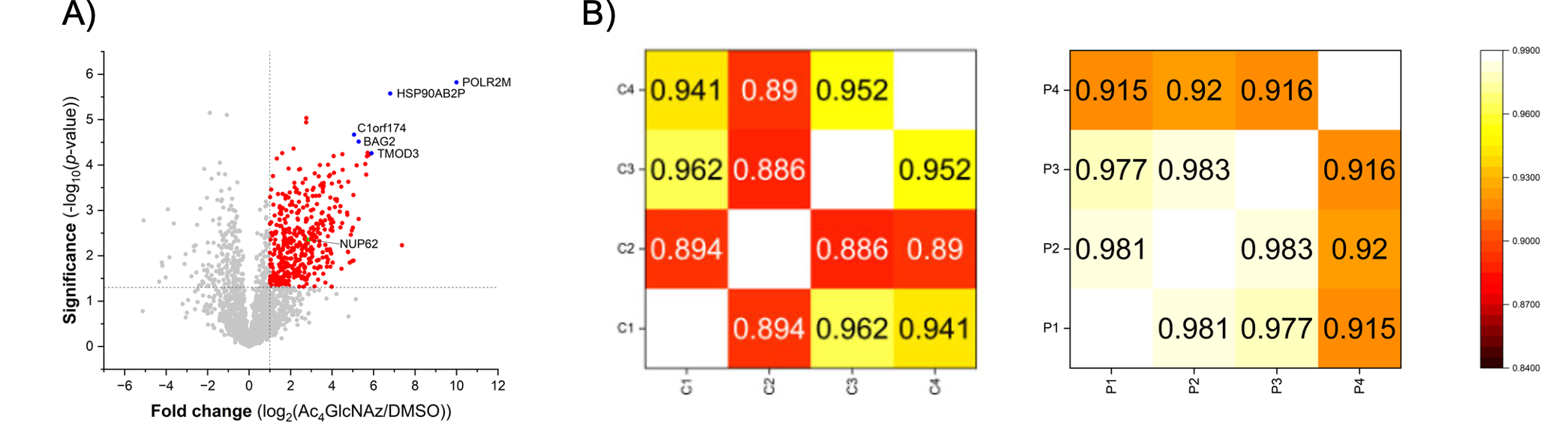

**Figure S5.** Volcano plot depicting the enrichment of *O*-GlcNAcylated proteins in HeLa cells. A) Significantly enriched proteins are shown as red circles, with the benchmark protein NUP62 highlighted in green. The five most significantly enriched proteins are indicated by blue circles (n = 4, cut-off p-value < 0.05, and fold-change > 1). B) Heatmap illustrating Pearson correlations among samples within each group.

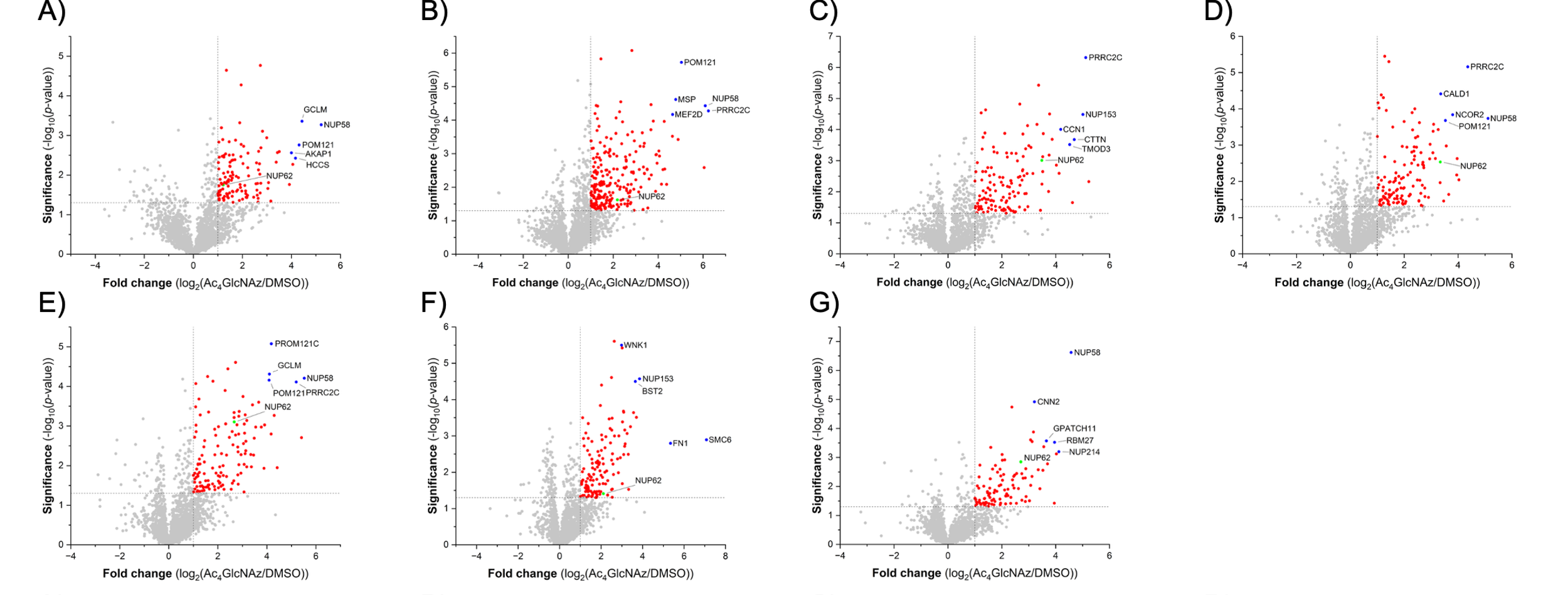

**Figure S6.** Volcano plots of the enriched *O*-GlcNAcylated proteins across a range of OSMI-1 concentrations in HeLa cells following the 25 µg SP2E workflow. Cells were treated with either a DMSO control (A) or increasing concentrations of OSMI-1: 10 nM (B), 50 nM (C), 100 nM (D), 500 nM (E), 1000 nM (F), and 2000 nM (G). The benchmark protein NUP62 is highlighted in green, and the top five most significantly enriched proteins are designated by blue circles. All significantly enriched proteins are shown as red circles. (n = 4, cut-off p-value < 0.05, and fold-change > 1).

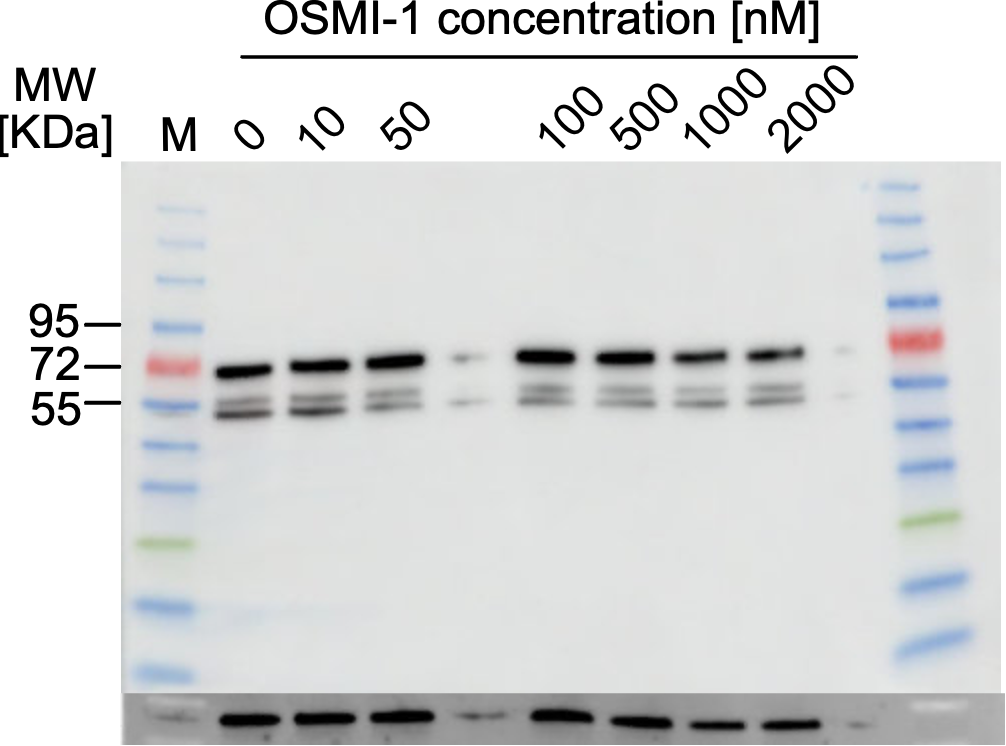

**Figure S7.** Western blot validation of NUP62 expression. HeLa cell lysates treated with the indicated concentrations of OSMI-1 (corresponding to Figure S5) were analyzed by Western blotting using an anti-NUP62 primary antibody. GAPDH was utilized as a loading control.

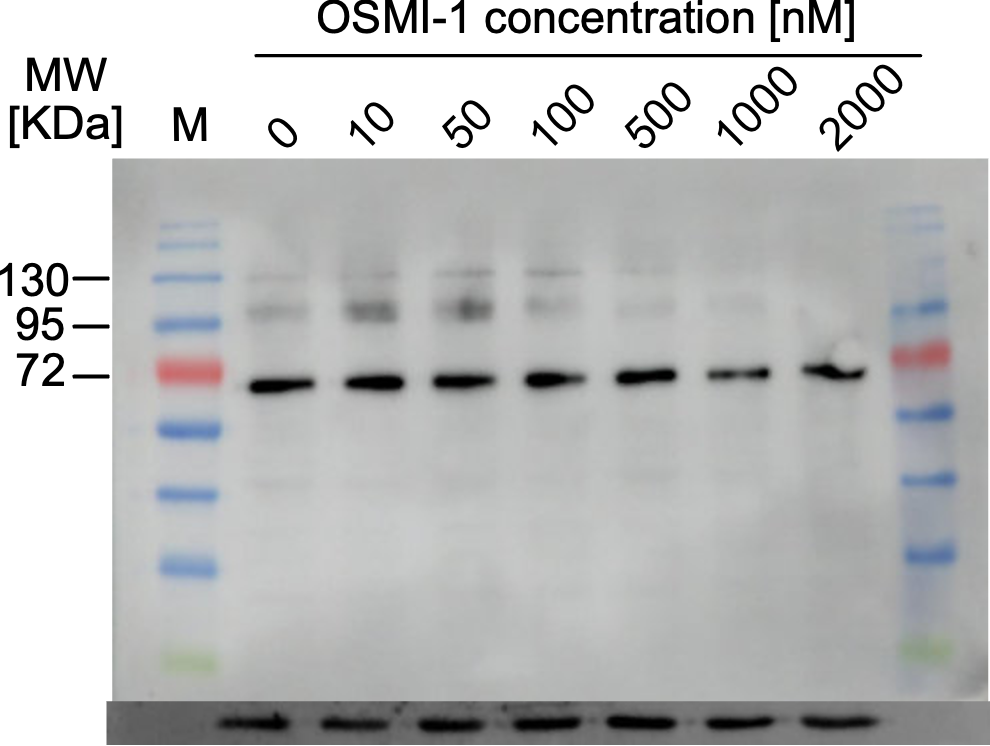

**Figure S8.** Western blot of global O-GlcNAcylation profiles. Representative immunoblot of HeLa cell lysates treated with varying concentrations of OSMI-1, matching the cohorts described in Figure S5. Primary detection was performed using an RL2 antibody to visualize global O-GlcNAcylated proteins. Lane loading equivalence was verified via GAPDH expression.

**MATERIALS AND METHODS**

**Key Resources Table**

| **Reagents or Resources** | **Source** | **Identifier** |
| --- | --- | --- |
| DMEM-high glucose | Sigma-Aldrich | Cat# D6546 |
| DPBS (1x) | Sigma-Aldrich | Cat# D8357 |
| Trypan Blue | Thermo Fisher Scientific | Cat# 11538886 |
| TrypLE Express | Thermo Fisher Scientific | Cat# 12604013 |
| Alanyl-Glutamine | Sigma-Aldrich | Cat# G8541 |
| FBS (Fetal Bovine Serum) | Thermo Fisher Scientic | Cat# A3840001 |
| Pro-N6pA | Santiago | CAS# 3004761-03-6 |
| Ac_4_GlcNAz | Jena Bioscience | Cat# CLK-1085-5 |
| Carboxylate coated magnetic beads (hydrophobic) | Cytiva | Cat# 65152105050350 |
| Carboxylate-coated magnetic beads (hydrophilic) | Cytiva | Cat# 45152105050350 |
| Streptavidin magnetic beads | New England BioLabs | Cat# S1420S |
| BSA | AppliChem | Cat# A6588 |
| HEPES  (*N*-2-Hydroxyethylpiperazine-*N*'-2-ethane sulphonic acid) | Carl Roth | Cat# HN77.5 |
| NP40 | Sigma-Aldrich | Cat# 74385 |
| SDS  (Sodium dodecyl sulfate) | AppliChem | Cat# A2572 |
| Biotin-PEG_3_-N_3_ | Carbosynth | Cat# FA34890 |
| DBCO-PEG_4_-Biotin | Jena Bioscience | Cat# CLK-A105P4-10 |
| 5/6-TAMRA-Azide-Biotin | Jena Bioscience | Lot# CLK -1048-5 |
| TBTA  Tris((1-benzyl-4 triazolyl)methyl)amine | TCI | Cat# T2993 |
| TCEP  tris(2-carboxyethyl) phosphine | Carbosynth | Cat# FT01756 |
| CuSO_4_ · 5H_2_O | Acros | Cat# 10627162 |
| Iodoacetamide | Merck | Cat# I1149-5G |
| H_2_O (LC-MS grade) | Carl Roth | Cat# AE72.3 |
| Urea | AppliChem | Cat# A1049 |
| Ethanol (EtOH) | Merck | Cat# 34852 |
| Acetonitrile (LC-MS grade) | Thermo Fisher Scientific | Cat# A955-212 |
| APS (Ammoniumperoxodisulfat) | Sigma-Aldrich | Cat# 09913 |
| Rotiphorese^TM^ Gel 30 (37,5:1) | Carl Roth | Cat# 3029.1 |
| TEMED  (N,N,N′,N′-Tetramethylethylenediamine) | Sigma-Aldrich | Cat# T9281 |
| Tris-base | Thermo Fisher Scientific | Cat# 10724344 |
| Glycine | Sigma-Aldrich | Cat# G8898-1KG |
| BenchMark™ Fluorescent Protein Standard | Invitrogen | Cat# LC5928 |
| TEAB  (Triethylammonium bicarbonate buffer) | Sigma-Aldrich | Cat# T7408 |
| Trypsin | Promega | Cat# V5113 |
| Formic Acid (LC-MS grade) | Thermo Fisher Scientific | Cat# A117 |
| Anti-O-Linked N-Acetylglucosamine antibody [RL2] | Abcam | Cat# ab2739 |
| Purified Mouse Anti-Nucleoporin p62 | BD Biosciences | Cat# 610498 |
| Amersham™ ECL Detection Reagents, | Cytiva | Cat# RPN2232 |
| Anti-mouse IgG, HRP-linked Antibody | Cell Signaling | Cat# 7076 |
| hFAB™ Rhodamine Anti-GAPDH Primary Antibody | Bio-Rad | Cat# 12004167 |
| OSMI-1 | Sigma-Aldrich | Cat# SML1621 |

**Buffers**

| **Buffer** | **Components** |
| --- | --- |
| Culture medium | DMEM(1x), 10% (v/v) FBS, 1% (v/v) Alanyl-Glutamine |
| Laemmli buffer (5 x) | 10% (w/v) SDS, 50% (v/v) glycerol, 25% (v/v) 2-mercaptoethanol, 0.5% (w/v) bromphenol blue, 315 mM Tris/HCl, pH 6.8 |
| Running buffer (10x) | 250 mM Tris, 1.92 M glycine, 1% (m/v) SDS |
| Semi-dry transfer buffer | 48 mM Tris, 39 mM glycine, 0.0375% (m/v) SDS, 20% (v/v) methanol |
